# Synaptic pruning, myelination and the emergence of psychiatric disorders in late adolescence

**DOI:** 10.64898/2026.05.20.726636

**Authors:** Bruno B. Averbeck, Nicolas Brunel

## Abstract

Adolescence is an important developmental period during which there are diverse changes in the brain and behavior. Goal-directed behaviors, including working memory, response inhibition, and reinforcement learning, improve during adolescence. At the same time there is substantial pruning of excitatory connections in prefrontal cortex and ongoing myelination of axons, which likely underlie these improvements in behavior. However, psychiatric disorders also become increasingly prevalent in late adolescence. In this study, we develop computational models that suggest a hypothesis for how ongoing changes in the brain can give rise to the increased prevalence of psychiatric disorders including depression, obsessive-compulsive disorder (OCD), schizophrenia and substance use disorders (SUDs).

We show that both myelination and pruning during adolescence lead to attractor landscapes in which strongly encoded memories, embedded in patterns of synaptic connectivity, come to dominate brain activity. Pruning and myelination lead to large, strong dynamical attractors which, if they are related to aversive emotions, can drive intrusive thoughts and compulsions in OCD, rumination in depression, or if they are related to drug exposure, compulsive drug seeking in SUDs. Thus, we suggest that psychiatric disorders emerge when early experience embeds pathophysiological patterns of synaptic connectivity in neural circuits. The neural changes during adolescent development then cause these memories to inflexibly dominate the neural and behavioral landscape.

## Introduction

Adolescence is an important developmental period (Larsen and Luna 2018), during which adult levels of performance are reached on a number of behaviors, including executive function (Gee, Bath et al. 2018, Wilbrecht and Davidow 2024). In parallel with these behavioral changes, there are ongoing changes in the brain, that underlie these improvements in behavior (Keshavan, Giedd et al. 2014, Larsen and Luna 2018, Averbeck 2022). Late adolescence, however, is also a period during which there is an increase in the emergence of psychiatric disorders (Paus, Keshavan et al. 2008, Uhlhaas, Davey et al. 2023). There are, at present, few quantitative hypotheses that link changes in the brain to changes in behavior, and the increased onset of psychiatric disorders.

Multiple forms of executive function including goal-directed behavior (Welsh, Pennington et al. 1991), working memory (Fry and Hale 2000, Gathercole, Pickering et al. 2004, Montez, Calabro et al. 2019) response inhibition (Liuzzi, Pine et al. 2023) and reinforcement learning (RL) (Palminteri, Kilford et al. 2016, Nussenbaum and Hartley 2019, Xia, Master et al. 2021) improve during adolescence. At the same time, ongoing neurodevelopmental processes continue to shape brain function, including synaptic pruning in the neocortex (Huttenlocher 1979, Petanjek, Judas et al. 2011) and myelination of long-range axons (Yakovlev and Lecours 1967, Miller, Duka et al. 2012, Grydeland, Walhovd et al. 2013).

In early development, neural systems establish connectivity with an abundance of synapses (Rakic, Bourgeois et al. 1986, Hensch 2004), which are subsequently pruned during adolescence through activity dependent mechanisms to establish adult connectivity and function (Huttenlocher 1979, Huttenlocher and Dabholkar 1997, Woo, Pucak et al. 1997, Kolluri, Sun et al. 2005, Petanjek, Judas et al. 2011, Watanabe and Kano 2011, Lee, Brott et al. 2014, Faust, Gunner et al. 2021). The number of inhibitory synapses and the number of neurons, on the other hand, remains relatively constant over development (Huttenlocher 1979, Bourgeois, Goldman-Rakic et al. 1994).

There is also ongoing myelination of long-range connections during adolescence including corticocortical, thalamocortical, and corticostriatal axons (Yakovlev and Lecours 1967). The largest increase in myelination occurs before the age of 5 (Yeung, Zdunek et al. 2014), even in association areas (Grydeland, Walhovd et al. 2013, Baum, Flournoy et al. 2022). However, myelination increases through adolescence and adulthood. Myelin increases conduction velocity and decreases the variability of synaptic delays (Pelletier and Pare 2002, Salami, Itami et al. 2003, Seidl 2014, Kato, Wake et al. 2020, Lefebvre, Clappison et al. 2025). Reducing variability in spike timing allows circuits to synchronize dynamic activity across nodes (Brunel 2000).

Although adolescence is an important developmental period for improvement on many behaviors, the emergence of multiple psychiatric disorders including obsessive-compulsive disorder (OCD), depression, schizophrenia, and substance use disorders (SUDs), peaks in late adolescence (Bethlehem, Seidlitz et al. 2022, Uhlhaas, Davey et al. 2023). The timing of this emergence suggests that changes taking place in the brain during this period increase the risk of developing a disorder (Paus, Keshavan et al. 2008). Thus, the same changes that improve executive function can lead to the development of psychiatric disorders.

Psychiatric disorders are characterized by inflexible, maladaptive behaviors including delusions in schizophrenia, drug seeking in SUDs, intrusive thoughts and compulsive rituals in OCD, and negative mood and rumination in depression. These behaviors have been conceptualized as dynamical attractors (Durstewitz, Huys et al. 2021, Yuan, Quan et al. 2026), and it has been suggested that an understanding of dynamics will be key for understanding psychiatric disorders (Rust 2025). For example, intrusive thoughts or ruminations are sequences of activity states in the brain toward which thoughts are strongly drawn, unintentionally, and despite efforts to avoid them. In this manuscript we explore the hypothesis that pruning and myelination can lead to the emergence of pathophysiological dynamical attractors that dominate behavior, when early life experience embeds strong synaptic connections that represent aversive emotional states or, in the case of SUDs, appetitive memories related to drug exposure. Thus, we assume that maladaptive attractors are present as a precondition, but adolescent development leads to the conditions under which they come to inflexibly dominate behavior.

## Results

Our goal in this study was to use computational modeling to understand how pruning and myelination, both important developmental processes that lead to improvements in cognition, may increase the risk of developing psychiatric disorders. We used reduced synaptic models, both rate and spiking networks, to characterize the conditions under which normal developmental processes that occur in adolescence can lead to the emergence of clinically distressful behavior. These models allow exploration of the effects of pruning and myelination, as these are important developmental changes that take place during adolescence.

Extensive work has shown that behavior, whether it is adaptive or maladaptive, is driven by patterns of neural activity that evolve in time (Fig. 1A-B)(Zhou, Seay et al. 2023, Tang, Bartolo et al. 2024). Maladaptive patterns of behavior are therefore governed by maladaptive patterns of neural activity. Therefore, we developed networks that learned sequences rather than fixed-points (i.e. stable patterns of population activity), because fixed points are not good models for dynamical neural activity underlying the behaviors we consider. Dynamical patterns of activity likely underlie many behaviors relevant to psychiatric disorders including cue induced craving in addiction, intrusive thoughts and compulsive rituals in OCD, delusional ideation in schizophrenia, and rumination in depression. Each of these represents a process that unfolds in time. Our networks learn to reproduce sequential patterns of activity (Figure 1C) leading to smooth dynamics in low-dimensional projections (Fig. 1D). Thus, they produce dynamical activity similar to that seen in the brain (Giarrocco and Averbeck 2026).

**Fig. 1.**
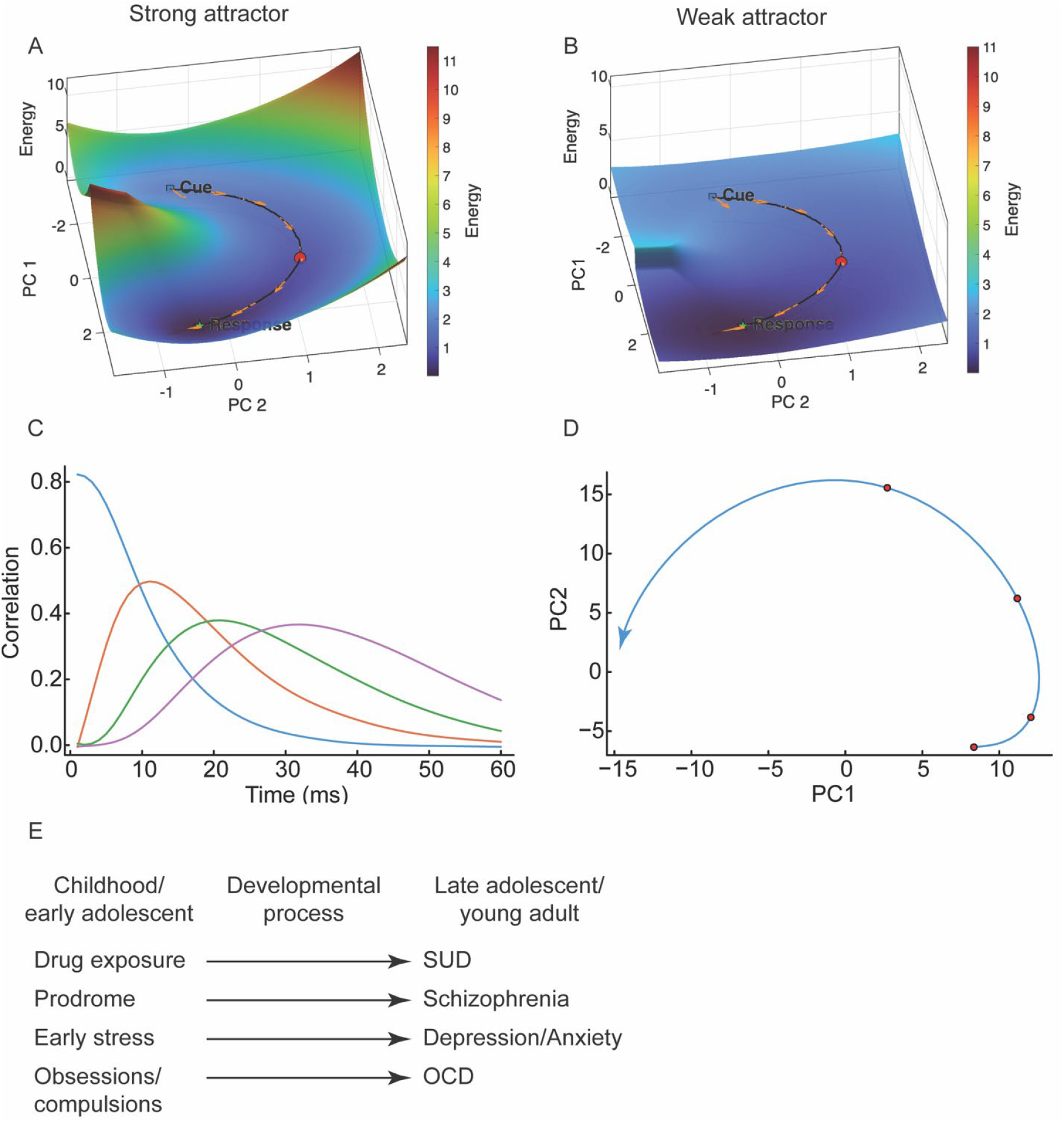
Attractors, sequences and hypothesis **A**. Activity in networks evolves from high to low energy. In this carton example, there is a drop in energy from the cue location to the response location, and the trajectory follows this energy gradient along the floor of the valley. Because the potential energy increases rapidly for directions that do not follow the trajectory (i.e. orthogonal to the direction of the trajectory), it is difficult to move activity in these directions. Therefore, this system would represent a strong attractor. **B**. Weaker attractor dynamics are represented by less steep energy landscapes. Two networks can produce the same average trajectory within different energy landscapes. It is easier to perturb activity off the average trajectory when the energy landscape is flatter. **C**. Example short sequence produced by a trained network. The correlation on the y-axis is the correlation coefficient between the network activity and each pattern in the sequence, where each pattern is a separate curve. The ordinal sequence of peaks represents the network progression through the sequence. **D**. Projection of network activity underlying the sequence shown in panel C onto a low-dimensional manifold defined by the first two principal components of network activity. Red dots indicate points where each sequence element peaks. **E**. Early experience can embed strong synaptic connectivity in networks. However, before pruning, these experiences are represented by dynamic activity with weak attractor dynamics. After pruning and myelination, however, attractor dynamics are strengthened. Particularly for early memories that were more strongly encoded.

Because behaviors are clinically distressful when they are inflexible and therefore difficult to suppress or change (for example compulsive drug seeking or compulsive rituals in OCD), we assume they come about due to strong attractor dynamics in the brain (Fig 1A-B). A strong attractor is an attractor that has high energy barriers around the trajectories it produces, where the energy represents the effort necessary to move activity in a particular direction in activity space. Strong attractors also have large basins of attraction, meaning that a large set of initial configurations tend to drive the network to the corresponding dynamical trajectory. Activity naturally evolves from higher to lower energy levels. However, if one wants to move activity from a lower to a higher energy level, effort must be exerted. Therefore, it is difficult to perturb activity away from trajectories surrounded by high energy barriers. Weaker attractors can produce similar trajectories (Fig. 1B). However, activity can be perturbed from its mean trajectory more easily.

The hypothesis we examine is that emotionally intense events in childhood and early adolescence embed strong synaptic connectivity in neural circuits (Fig. 1E). In most cases of relevance to psychiatric disorders these would be aversive events but in the case of substance use disorders they would be euphoric experiences. For example, it is known that early exposure to drugs of abuse, which drives strong synaptic changes in the brain (Kauer and Malenka 2007), leads to an increased lifetime risk of addiction (Anthony and Petronis 1995). Similarly early life stress leads to increased risk of later depression and anxiety (Wade, Wright et al. 2022), and prodromal periods in schizophrenia (White, Anjum et al. 2006) may embed strong delusional associations in synaptic connectivity, perhaps due to aberrant dopamine activity (Kapur, Mizrahi et al. 2005). Furthermore, a significant fraction of patients with OCD identify traumatic events as precipitating their compulsive behavior (Pinciotti and Fisher 2022, Pinciotti, Thorsson et al. 2026). These early events drive synaptic plasticity and affect behavior. However, before adolescent development these behaviors are less rigid and they do not lead to clinical levels of behavioral pathology. Subsequently, during adolescent development when pruning and myelination take place, these relatively strong synaptic connections and their associated neural dynamics come to dominate the attractor landscape leading to inflexible behaviors. When these brain dynamics are related to maladaptive behaviors, they lead to psychiatric disorders. The same developmental processes can, however, improve the neural computations that underlie executive functions when they refine synaptic connectivity related to cognitive processes (Averbeck 2022). Thus, the same developmental process that strengthens attractor dynamics underlying cognitive operations and therefore makes executive function more reliable and less susceptible to distractors, can make maladaptive behaviors inflexible and rigid.

In the results that follow, we first confirm that networks can be trained with Hebbian plasticity rules to generate sequential patterns of activity (Murray and Escola 2017, Gillett, Pereira et al. 2020, Gillett and Brunel 2024). We then show how these patterns can model Pavlovian associative behavior as a model of fear conditioning, extinction, and reacquisition. Because dynamical patterns are ubiquitous in the brain, our results are more general and likely represent processes like rumination in depression and the execution of rituals in OCD. We then examine the effects of pruning and myelination on attractor dynamics. Our results show that pruning and myelination preferentially strengthen dynamics associated with sequences that are strongly encoded, i.e. encoded with stronger synapses, relative to sequences that are weakly encoded. Thus, pruning and myelination preferentially strengthens attractors associated with strongly encoded memories.

We began by examining the coding of sequences of activity in networks trained with Hebbian plasticity rules. Exposing the network to sequential patterns of activity leads to synaptic connectivity changes that encode the sequence. When networks were trained incrementally, synaptic connectivity related to the sequence was gradually strengthened. The networks were then probed for their ability to generate the sequences by initializing them with the first pattern and then letting them run autonomously. As the synaptic connectivity strengthened, they acquired the ability to autonomously generate the sequences (Fig. 2A). Overlapping sets of single neurons were activated as the sequences unfolded (Fig. 2B-C). Note that previous work has shown that the speed of sequence retrieval is controlled by the time constant of rate dynamics (Gillett et al 2020) and the balance between temporally symmetric and temporally asymmetric plasticity (Gillett and Brunel 2024). Here, we chose for simplicity to use parameters similar to Gillett et al 2020, leading to fast sequence retrieval (∼200ms). Much slower retrieval could be obtained by introducing a temporally symmetric term in synaptic connectivity, as shown by Gillett et al (2024).

**Fig. 2.**
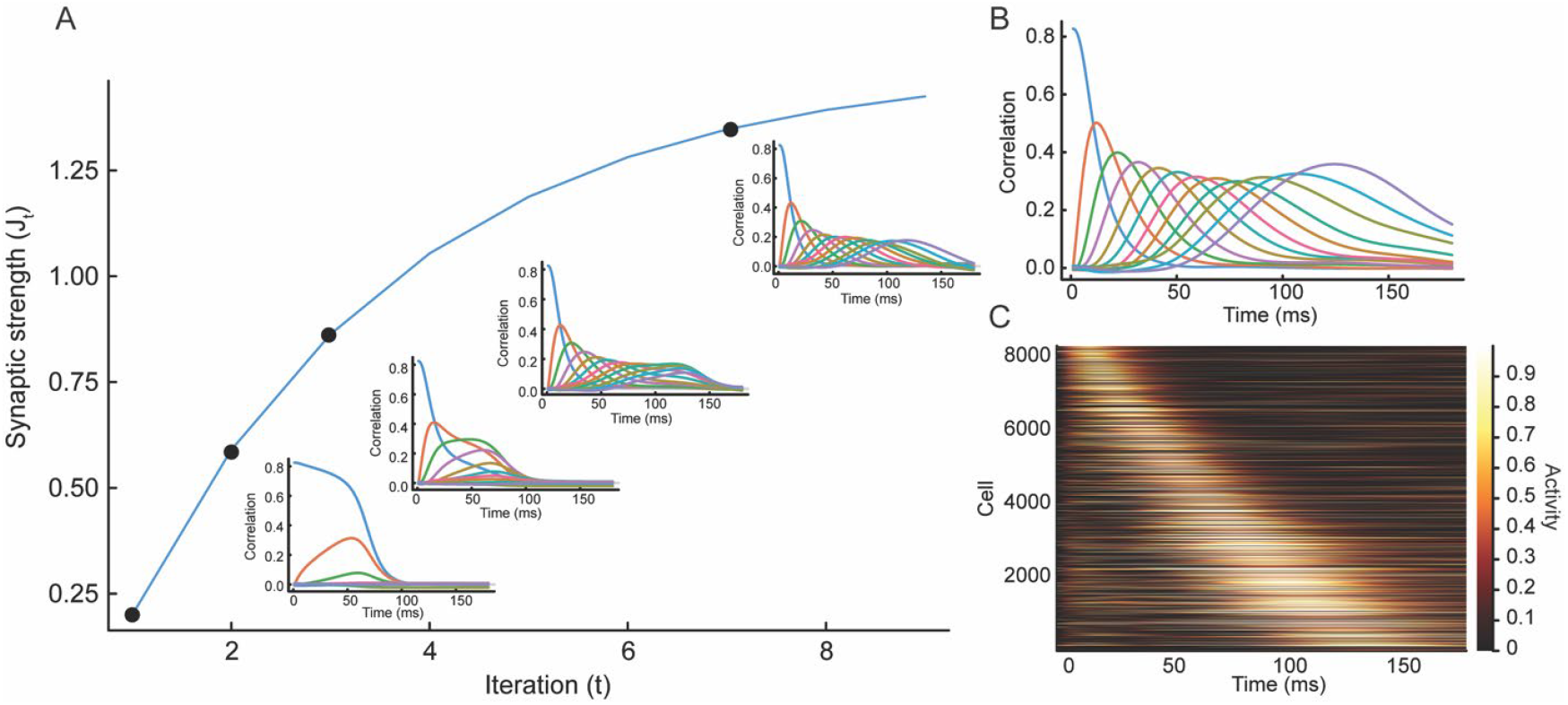
Sequence learning and example activity. **A**. Sequence learning iterations vs. synaptic strength, and example sequence recall. Network connectivity was trained by exposing the network to the sequence and updating the synaptic connectivity with spike-timing dependent plasticity. Each training iteration strengthened the synaptic connectivity related to the sequence. The ability of the network to generate the sequence autonomously was then probed by starting the network with the first pattern of activity and then letting the network run autonomously. Black dots indicate points at which examples are sampled. The pattern of correlations between network population activity and sequential patterns, with overlaps between adjacent sequence elements, are similar to those measured in prefrontal cortex population activity in the brain (Averbeck, Chafee et al. 2002). For simulation parameters, *J*^*s*^ = 1.5, *σ*_*r*_ = 0. **B**. Sequence recall (synaptic strength = 1.1). **C**. Activity of 8300 example neurons, sorted by time of maximum activity.

Next we examined this as a model for learning and extinction of aversive associations. Although many models can be used to capture these learning processes, our goal was to establish that our learning mechanism can capture fundamental features of these learning processes, before examining the effects of pruning and myelination for which our model is well suited. For disorders including anxiety and OCD, extinction training is an effective treatment. During extinction training in anxiety disorders, a cue that ostensibly predicts a negative outcome is followed by no outcome. In OCD presentation of a cue that normally would drive execution of a ritual is followed by inhibition of the ritual. Therefore, we next explored the ability of the networks to model Pavlovian conditioning, extinction, and reacquisition, as an example of these processes. Networks were first trained to associate a cue with an aversive outcome (Fig. 3A). (Note that our states are arbitrary patterns of population activity. However, patterns of activity in the amygdala, for example, represent appetitive and aversive states (Tang, Bartolo et al. 2024).) The cue was modeled as the first three patterns of activity, and the aversive outcome as the remaining patterns in the sequence, similar to the situation in which an aversive outcome is presented at cue termination during learning. Exposing the network to patterns related to the cue and outcome led the networks to acquire the Pavlovian association (Fig. 3D), where the association is reflected in the network autonomously generating a sequence that links cue states (the first three sequence elements) to negative association states (the later elements of the sequence). When we examined the correlation between activity in the network, and the last pattern in the sequence, reflecting the learned association, we found that this correlation quickly increased during initial acquisition (Fig. 3B). Further, when we used network activity to predict avoidance behavior, by decoding the last element of the sequence, we found that it quickly increased during acquisition, reaching a peak after only a few exposures (iterations; Fig. 3C).

**Fig. 3.**
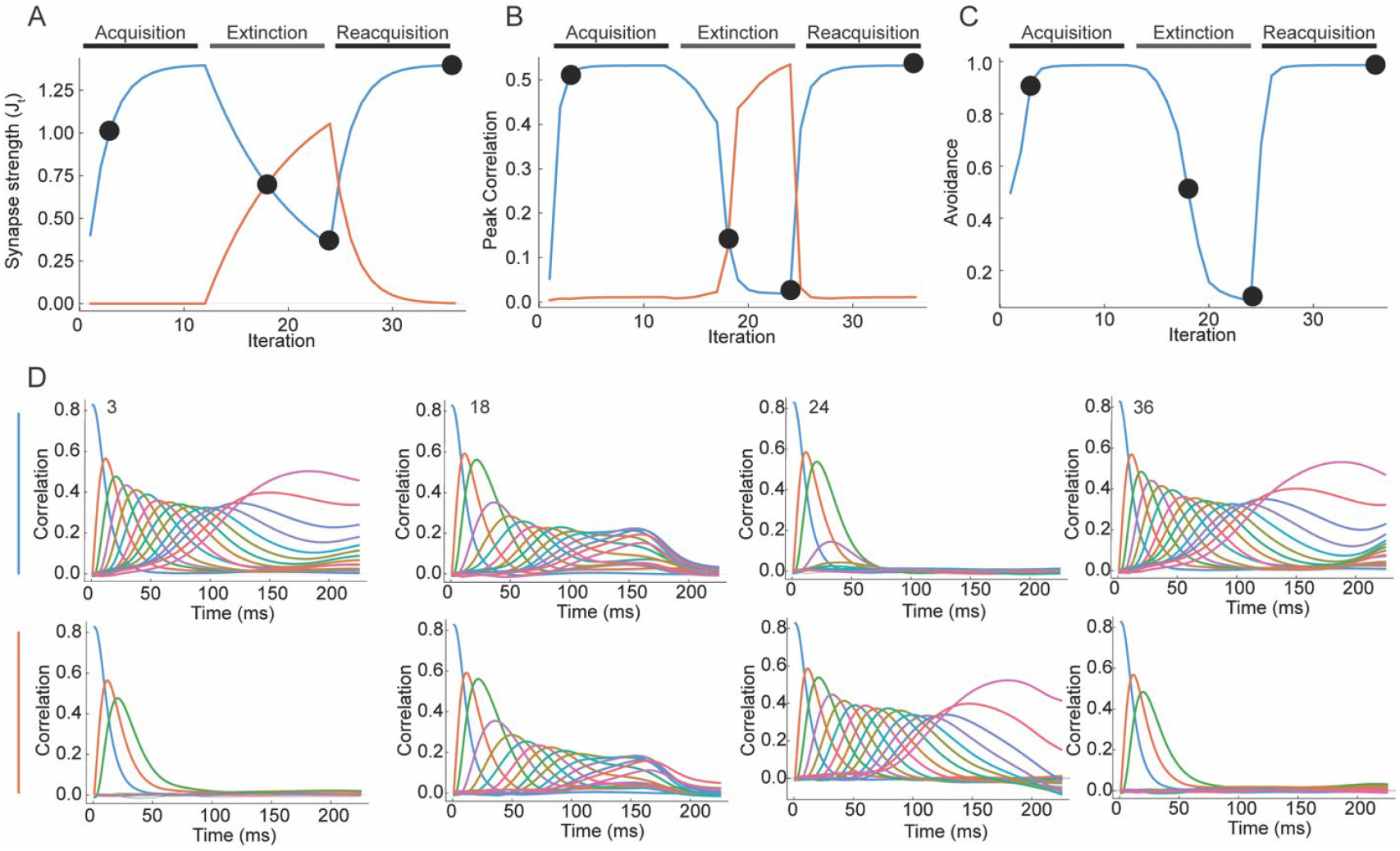
Acquisition, extinction, and reacquisition of sequential activity. For simulation parameters, *J*^*s*^ = 1.4, *σ*_*r*_ = 0. During acquisition and reacquisition of a fear memory, *ρ* = 0.4. During extinction (i.e. blue curve during extinction and red curve during reacquisition), *ρ* = 0.11. **A**. Acquisition of sequence representing Pavlovian association of cue and aversive outcome. Blue curves are association of cue (first three sequence elements) with aversive outcome (final element). Red curves are association of the same cue with no outcome. 1-12 are initial acquisition, 13-24 are extinction, and 25-36 are reacquisition. Black dots indicate points at which sequence recall examples are shown in Panel D. **B**. Peak correlation with the last sequence element during sequence recall. Blue curve indicates correlation with aversive association, red curve indicates correlation with safety association. **C**. Avoidance behavior, decoded from population neural activity in the last sequence element representing either predicted aversive or no outcomes. **D**. Example sequence recall taken from points indicated by black dots in panels A-C. Top row, are for aversive associations, and bottom row are for safety associations. Note that the first three patterns correspond to the cue and are the same in both rows. The inset numbers correspond to the black dots indicated in panels A-C.

Next, the network underwent extinction training. It is known from behavioral work that extinction learning does not erase the aversive memory (Bouton, Maren et al. 2021). Rather a new memory is learned, representing the lack of an aversive outcome, or no outcome (Macdonald, Ma et al. 2025). During extinction training, the network was exposed to the same cue, but the cue was not followed by an aversive US. Rather the cue was followed by no outcome. At the same time, synaptic connectivity related to the aversive outcome association slowly decayed (Fig. 3A). When the network was run autonomously, and the last pattern of activity generated by the network was compared with the aversive and no outcome patterns, we found that the network shifted during extinction training to generate the pattern related to the no outcome, and not the aversive outcome (Fig. 3B, D). Similarly, when we predicted avoidance behavior, it was extinguished during extinction training (Fig. 3C).

During extinction the safety prediction came to dominate, and the network generated patterns of activity that predicted a neutral outcome. However, it could be seen that the underlying synaptic connectivity related to the aversive prediction had only partially decayed (Fig. 3A). Thus, when the network was re-exposed to aversive cue-outcome associations, it quickly began to generate patterns related to the aversive outcome (Fig. 3B, D). We then allowed the network to reacquire the aversive association, by running more trials with an aversive outcome, and found that the network quickly reacquired the aversive association.

### Three-factor learning, synaptic pruning and attractor dynamics

Next, we examined the effects of three-factor learning on sequence representation. In several systems, it has been shown that neuromodulators, for example norepinephrine, affect Hebbian learning by gating or modulating plasticity (Fremaux and Gerstner 2015). When neuromodulators are at low concentrations there is no Hebbian learning, and when concentrations are increased, Hebbian learning is stronger. It is known as three-factor learning because it requires the coincidence of pre-synaptic activity, post-synaptic activity, and the appropriate neuromodulatory tone. We used three-factor rules to simulate sequence learning under different levels of emotional intensity, where the emotional intensity of the event drove different levels of neuromodulator release. More specifically, we trained networks on three sequences and implemented three-factor learning by modulating both the rate and asymptotic strength of learning of each sequence differentially (Fig. 4A). When we examined sequence recall in these networks, all three sequences could be recalled in this example under noise-free conditions, when the networks were started with the first pattern of each sequence (Fig. 4B-D). This shows that networks can be trained with multiple sequences, where each sequence has a different strength of representation driven by the interaction of the concentration of the neuromodulator and the Hebbian learning rule.

**Fig. 4.**
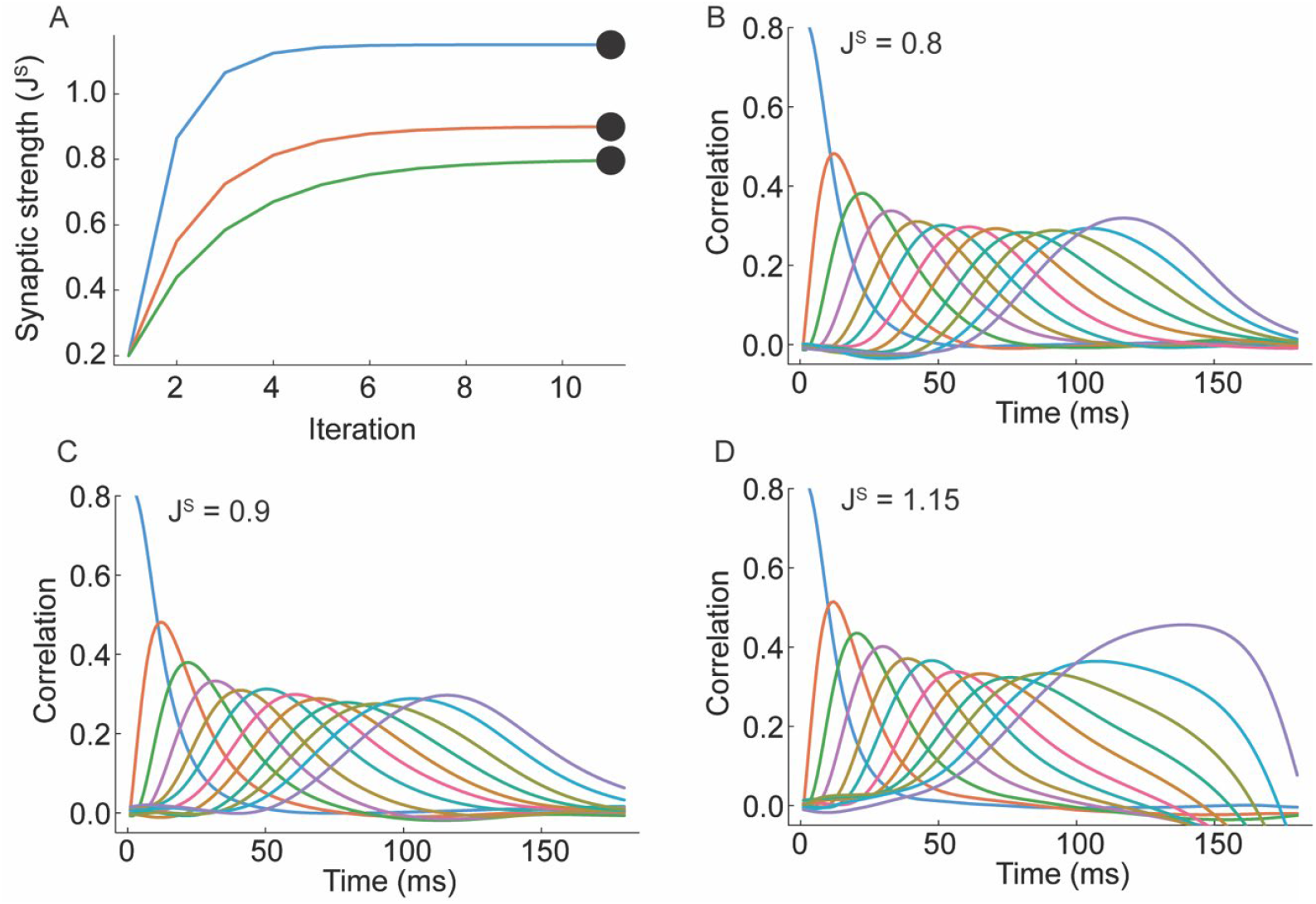
Three-factor learning and synaptic strength. **A**. Example simultaneous learning of three different sequences with different dopamine neuromodulation. Neuromodulation affects the rate of learning and the final strength of the sequence. **B**. Sequence recall of a sequence with final overall synaptic strength of J^S^ = 0.8, *ρ* = 0.4. **C**. Sequence recall of a sequence with final overall synaptic strength of J^S^ = 0.9, *ρ* = 0.5. **D**. Sequence recall with final overall synaptic strength of J^S^ = 1.15, *ρ* = 0.7, *σ*_*r*_ = 0.

Next, we examined the effect of synaptic pruning on recall and the size of attractor basins in networks trained with three sequences with different levels of final synaptic strength. Thus, networks first learned three sequences with different levels of neuromodulation (Fig. 4), with their full synaptic connectivity. We then pruned the weakest connections in these networks and used homeostatic synaptic scaling to strengthen the remaining synapses (Chung, Wills et al. 2017, Li, Ma et al. 2017) such that the total excitatory and inhibitory synaptic drive in the network was constant (Fig. 5E).

**Fig. 5.**
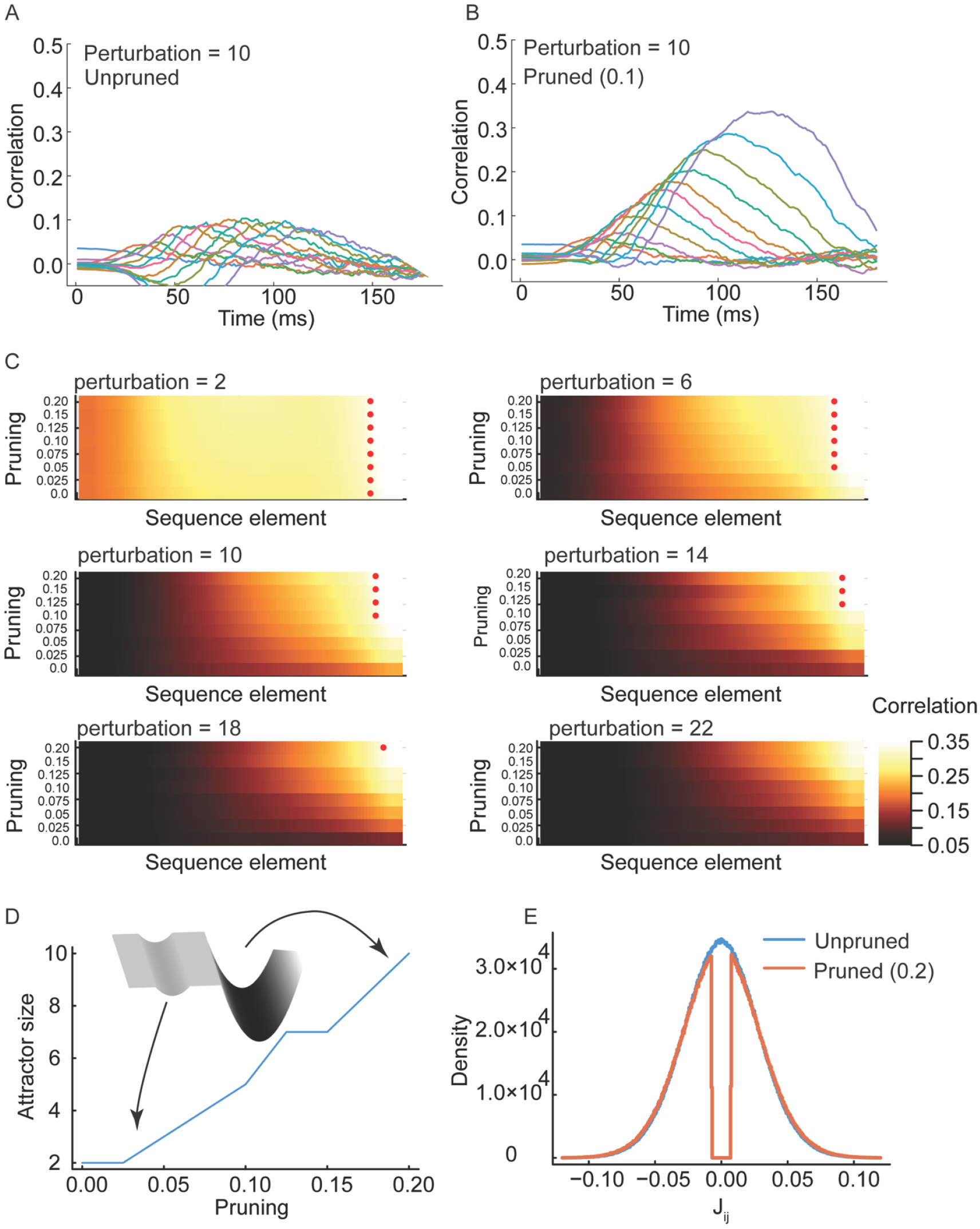
Sequence recall in pruned and unpruned networks with final overall synaptic strength of J^S^ = 1.15, *σ*_*r*_ = 0.3. **A**. Sequence recall in unpruned network when initial pattern of activity is perturbed by adding Gaussian white noise with a standard deviation of 6 to the first pattern. **B**. Recall of the same sequence in a network in which 10% of the weakest connections have been pruned, and the remaining connections have been homeostatically rescaled to maintain the same total strength of synaptic inputs. **C**. Averages across a series of perturbation strengths (*σ*_*p*_) and levels of pruning, indicated by the fraction of pruned synapses for each row. Correlation indicates the peak correlation of each element of the sequence. Plots are averages over N = 96 random networks and perturbations. Red dots indicate the points at which the recall exceeds 0.35. **D**. Pruning vs. attractor size. Attractor size was estimated as the maximum perturbation at which the final sequence element exceeded a correlation of 0.35. Note in general that increased pruning consistently increased the strength of the sequence recall across levels of perturbation. Insets are cartoons to indicate that increased pruning increases the size of the dynamical attractor. **E**. Example distribution of individual connection strengths, J_ij_. Not that the pruned curve has more strong connections, but the effect across the distribution is subtle.

We first examined sequence recall of the strongest sequence, with a final synaptic strength of J^S^ = 1.15. To examine the size of the attractor basin, we perturbed the initial pattern of activity with Gaussian white noise, with different standard deviations (Fig. 5C). Thus, rather than start the network perfectly aligned with the first pattern of activity in the sequence, we started the network with a pattern that deviated from the first pattern, with the size of the deviation given by the standard deviation of the white noise. If the network still recalled the sequence, even though it was not started aligned with the first element, the initial pattern was in the basin of attraction for that sequence. By examining whether the sequence could be recalled with different levels of noise, we could characterize the size of the basin of attraction for each sequence, for different levels of pruning. Networks that recalled sequences when started with a pattern that was far from the first pattern had large basins of attraction, and networks that failed to recall sequences when started far from the first pattern had smaller basins of attraction.

Sequence recall was weaker in unpruned networks (Fig. 5A) and more robust in pruned networks (Fig. 5B) when the initial activity was perturbed with noise. We summarized recall performance by assessing the peak correlation of each sequence element and plotting this as a function of pruning, averaged across 96 networks (Fig. 5C). Across a range of perturbations and levels of pruning, more pruning led to stronger recall at each noise level (Fig. 5C). Thus, pruned networks had larger basins of attraction. When we characterized whether recall of the last element exceeded 0.35 (as an arbitrary threshold to summarize the results) across noise levels, we found that the size of the basin of attraction (estimated as the noise level at which recall exceeded 0.35) increased as the networks were further pruned (Fig. 5D). Thus, pruning increased the basin of attraction for the strongest sequence.

When we examined sequence recall for the weaker sequences (Fig. 6), we found that, while pruning did increase the strength of recall for the weakest perturbations (Fig. 6A, B, left), sequence recall was not robust at stronger perturbations. Thus, there was minimal recall for even moderate levels of noise (Fig. 6A, B, right). The basins of attraction, therefore, for the weaker sequences were limited. Overall, pruning led to a substantial increase in the basin of attraction for the stronger sequences, such that activity could be started far from the first element, and the sequence was still recalled. Pruning did not lead to robust sequence recall, or expansion of the basin of attraction, for weaker sequences.

**Fig. 6.**
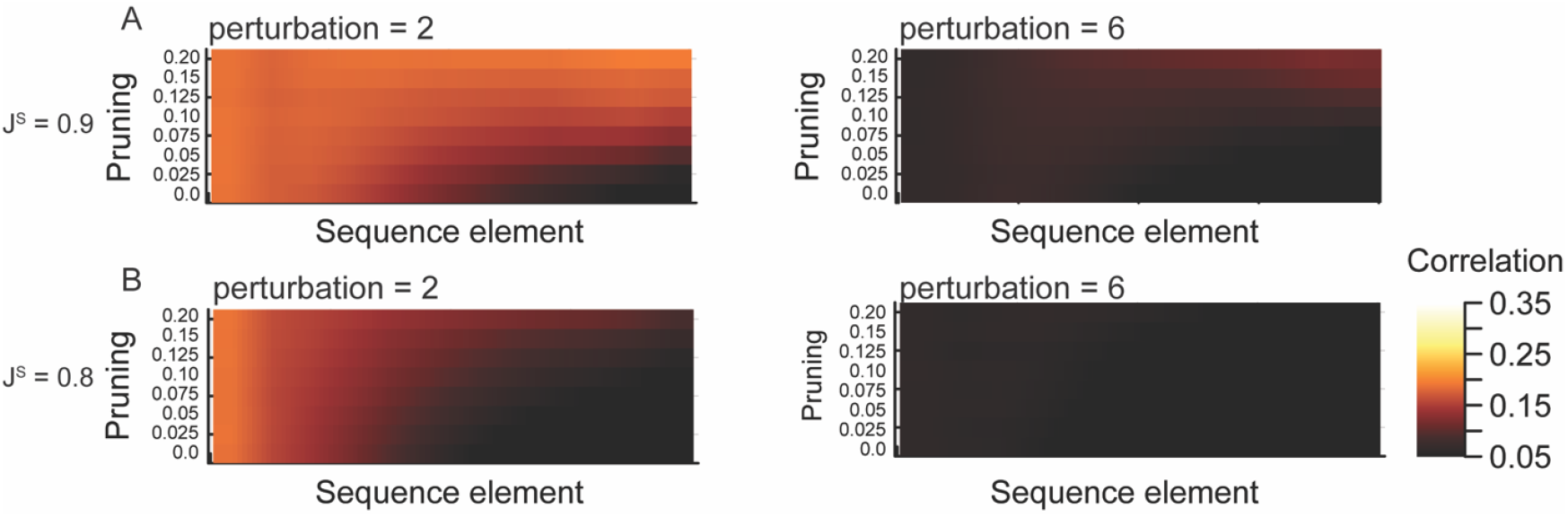
Average recall of sequences with weaker synaptic strength. Same parameters as Figure 4. **A**. Left panel shows recall with low perturbation and different levels of pruning for sequence with overall synaptic strength of J^S^ = 0.9. Right panel shows the same with higher perturbation. **B**. Same as A for sequence with overall strength of J^S^ = 0.8.

### Spiking networks, sequence recall and the effect of myelination

Next, we moved from rate networks to excitatory and inhibitory (E/I) spiking networks. E/I networks allow us to restrict pruning to excitatory connections, consistent with biological pruning in prefrontal cortex (Petanjek, Judas et al. 2011). They also allow us to model the effect of myelination on long-range cortical-cortical connections, because spike latency effects can be explicitly modeled.

We first examined sequence recall in a single network of E/I spiking neurons and replicated previous results that showed robust recall in these networks (Fig. 7). Next, we built a parietal-frontal network with two populations of neurons (Fig. 8). All inhibitory connections were restricted to the local population. Excitatory connections were both local (recurrent) and long-range. Long-range connections were bidirectional such that parietal cortex sent connections to prefrontal cortex, and prefrontal cortex sent return connections to parietal cortex. Local connectivity in all networks had 1 ms delays. Pruning in these networks (e.g. Fig. 8B) was restricted to excitatory connections within each population. Neither long-range connections nor inhibitory connections were pruned. The effect of myelination was modeled as changes in spike time delay distributions on long-range connections (e.g. Fig. 8C). Increasing myelination increases conduction velocity between areas. More relevant to our simulations, however, myelination decreases variability in synaptic delay distributions. Thus, we simulated long-range connections with several distributions of synaptic delays.

**Fig. 7.**
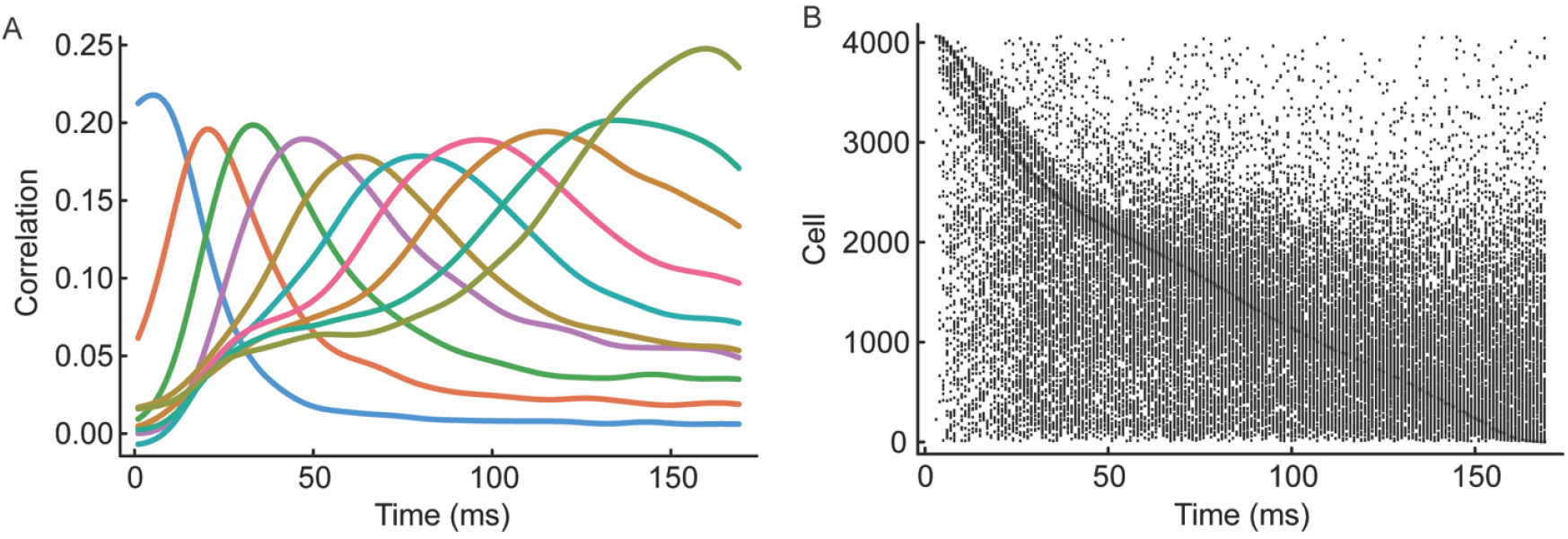
Sequence recall in a network of excitatory and inhibitory spiking neurons. **A**. Sequence recall in network. **B**. Activity of excitatory neurons in the network sorted by time of peak activity in each neuron.

**Fig. 8.**
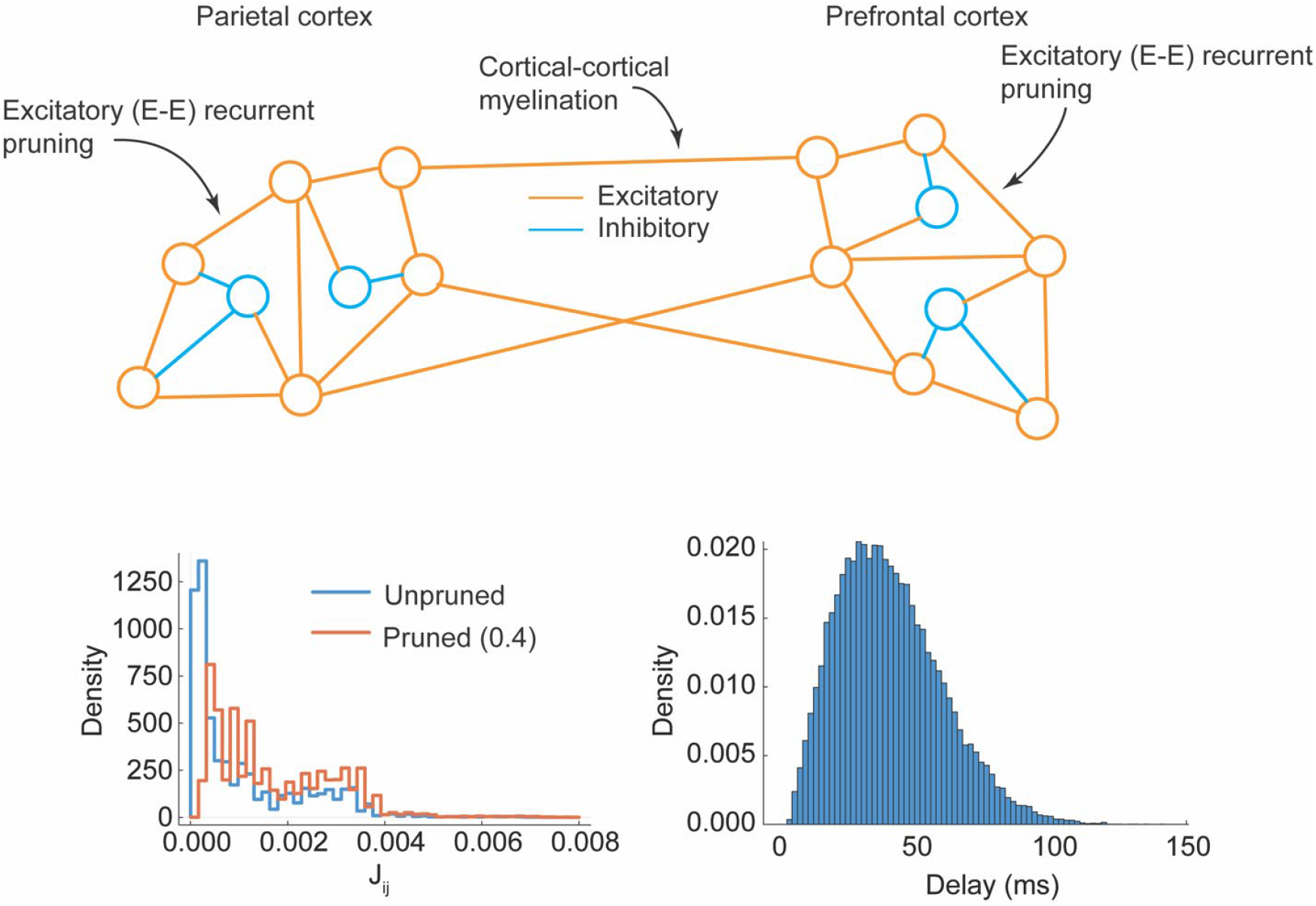
A. Network to test effects of pruning and myelination in connected populations of spiking neurons. Each area has N = 15000 excitatory neurons and N = 3750 inhibitory neurons. All inhibitory connections are local. Cortical-cortical connections are excitatory. Local circuit connections are assumed to be fast (i.e. delay = 1 ms). Myelination only affects the distribution of delays on cortical-cortical connections. Pruning was restricted to local E-E connections. E-E connections between populations were not pruned. E-I and I-E connections were not pruned. Weight distributions undergo homeostatic synaptic rescaling to maintain strength of excitatory inputs to each neuron. Note that cortical-cortical connections are bidirectional. Population 1 sends excitatory connections to population 2 and population 2 sends excitatory connections to population 1. Myelination affects delay distributions in both directions. **B**. Example distribution of local recurrent excitatory connections before and after pruning. **C**. Example distribution of delays for long-range excitatory connections, between populations, with location parameter = 30 ms.

We examined propagation of sequences across populations, by starting a sequence in only one population (e.g. parietal cortex), and examining whether the second population (e.g. prefrontal cortex) entrained to the first population (Fig. 9). When we compared sequence coupling between networks with broad synaptic delay distributions on long-range connections (mode = 30 ms, Fig. 8C) to networks with highly constrained, 1 ms delays on long-range connections, we found that coupling was more robust in myelinated networks with narrow delay distributions (Fig. 9A, 9B, right panel). Interestingly, because of the feedback connections from prefrontal to parietal cortex, representations in the networks in which the sequences originated (parietal cortex) also showed stronger sequence representations, even though connectivity and pruning was identical within local networks (Fig. 9A, 9B, left panel). The only difference between the networks was the distribution of delays on long-range connections. Local connections in both networks had the same synaptic connectivity, delays, and pruning. When we compared coupling across areas in these networks for a range of pruning levels, we found that coupling was consistently robust in the network with 1 ms delays, compared to the network with a broad delay distribution (Fig. 9C, 9D). Pruning, however, further strengthened coupling. Thus, both pruning and myelination increase sequence coupling across networks of neurons.

**Fig. 9.**
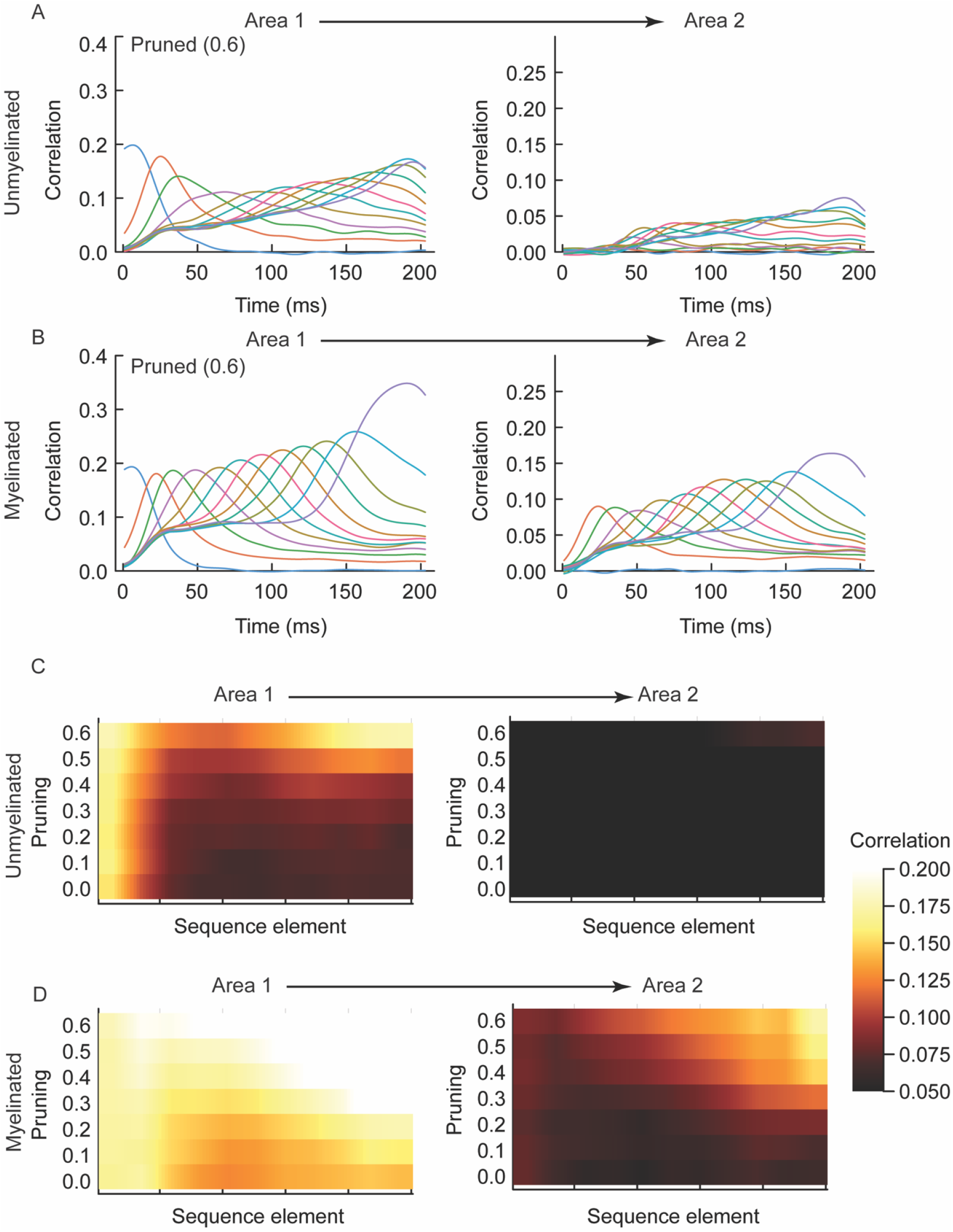
Sequence coupling. Plots are averages over 12 iterations. **A**. Average sequence recall in population 1 and 2 when the sequence activity is started in only population 1 with 30 ms modal average synaptic delay distribution. Sequence activity in population 2 has to propagate from population 1. Pruning = 0.6. Network activity was perturbed with *σ*_*p*_ = 6 in both populations. Note that arrow indicates propagation of sequence from population 1 to 2, but population 2 sends feedback connections to population 1. **B**. Same as C for 1 ms delays between all neurons, including long-range connections. Pruning = 0.6. **C**. Peak correlation of sequence elements in population 1 (left panel) and 2 (right panel) with 30 ms modal delay distribution for different levels of pruning. **D**. Same as panel C for 1 ms delays.

When we examined various cortical-cortical synaptic delay distributions (Fig. 10A), we found that sequence coupling to prefrontal cortex was consistently weaker as delay distributions became more variable (Fig. 10B). Thus, myelination leads to increased coupling of sequential activity across areas connected by long-range connections in which myelination decreases variability in delay distributions.

**Fig. 10.**
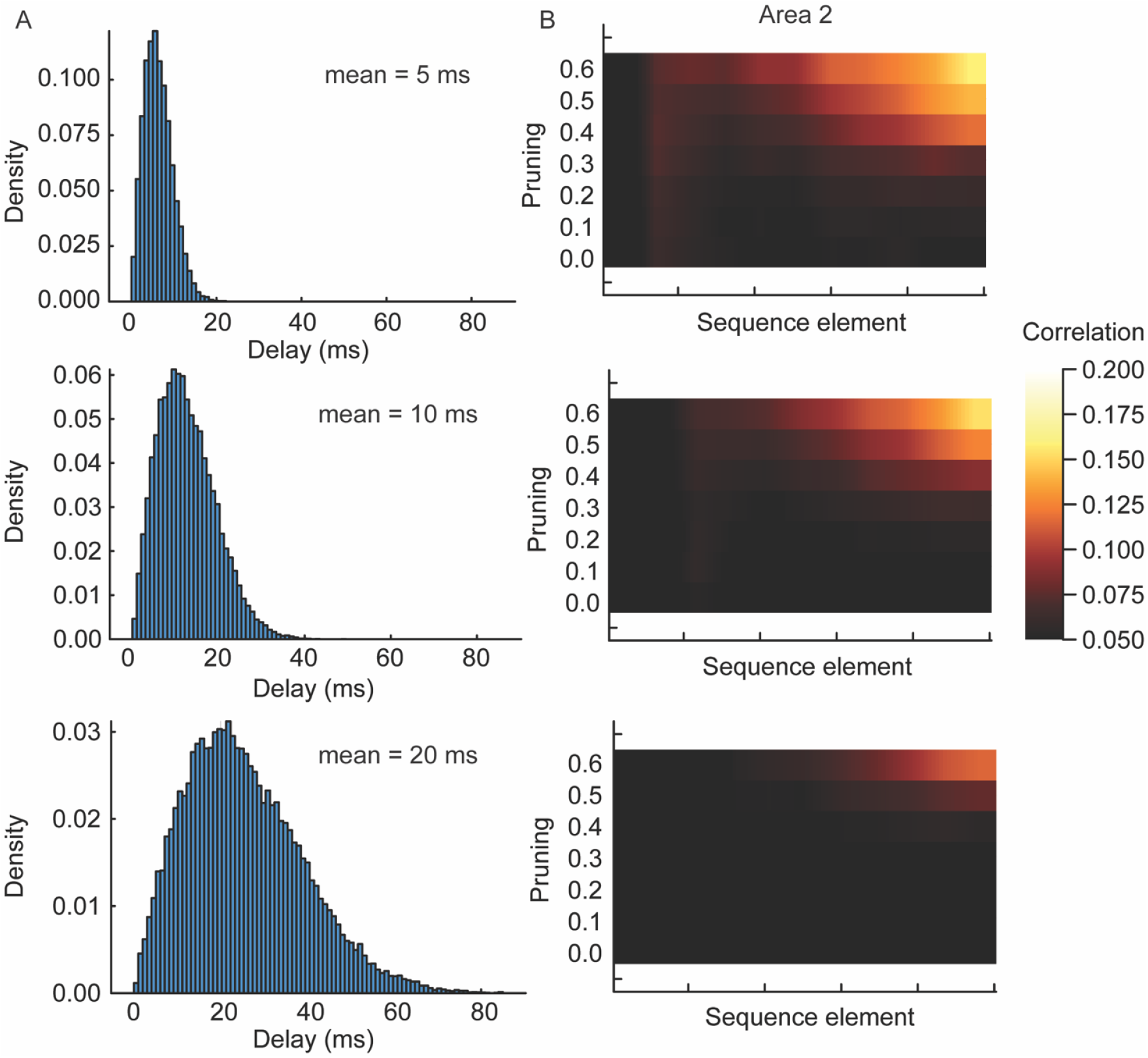
Average sequence recall in population 2 for different delay distributions. All other parameters are the same as Figure 9. **A**. Delay distributions. **B**. Average sequence recall in population 2 at different pruning levels, for each delay distribution.

## Discussion

Here we have shown that synaptic pruning in rate networks, and excitatory synaptic pruning and increased myelination in spiking networks lead to networks in which emotionally intense memories, with strong synaptic connectivity, can dominate attractor landscapes. While the fractional change in excitatory connections during adolescence (Petanjek, Judas et al. 2011) is larger than the fractional change in myelination (Baum, Flournoy et al. 2022) during the same period, both contribute to changing the attractor landscape. Emotionally intense memories create strong synaptic connectivity through three-factor learning rules (Reynolds and Wickens 2002, Pawlak, Wickens et al. 2010, Fremaux and Gerstner 2015). Before synaptic pruning, memories for sequences encoded with both strong and weak synapses can be recalled when they are triggered by a cue which is well-aligned with their initial elements.

Following pruning, however, activity patterns far from the initial patterns of sequences that were initially encoded with strong synaptic connectivity, drive recall of those sequences. Less strongly encoded sequences, however, are not well-recalled when activity patterns are not well-aligned. Thus, the attractor basin for strongly encoded sequences is expanded, at the expense of less-strongly encoded sequences, through synaptic pruning.

Myelination affects long-range connections including cortical-cortical (modeled here), thalamocortical, and corticostriatal axons. Myelination increases conduction velocity, which decreases synaptic delays between areas (Salami, Itami et al. 2003, Kato, Wake et al. 2020) and decreases the variability of delays between areas (Pelletier and Pare 2002, Seidl 2014). We found that cortical-cortical connections with narrow and short synaptic delay distributions led to stronger coupling of sequences in target areas, and stronger recall within areas in which sequence recall originated. Thus, increased myelination leads to connected networks across which sequential memories couple more effectively. While both pruning and myelination can contribute to the increase in executive function seen during adolescence, they can also unmask embedded emotionally intense memories, which can come to dominate attractor landscapes.

Previous modeling work has shown that synaptic pruning can lead to networks that are more robust to noise, when they are trained on working memory and reinforcement learning tasks (Averbeck 2022). This study showed that pruned networks trained on working memory tasks were resistant to distraction, modeled as a perturbation of neural activity, during delay periods. When these networks were perturbed away from mean trajectories that encoded working memory for cues during delay periods, they were pulled back towards the mean trajectories and accurately maintained items in memory. In addition, pruning these networks during training led to connectivity distributions like those obtained in the networks in the present study with few small connections (as these were pruned away) and more strong connections. These modeling results were followed-up by analyses of longitudinal EEG data, which attempted to characterize whether variability, which is an intrinsic perturbation, in single-trial evoked EEG data in adolescents also showed evidence for stronger attractor dynamics (Liuzzi, Pine et al. 2023, Andujar, Liuzzi et al. 2026). These studies found that variability in evoked activity decayed back to mean trajectories faster when participants carried out this task in their late (18 y.o.) vs. early (12 y.o) teens. While pruning may primarily drive the effects seen in the EEG data, myelination may also contribute, given the results of the present study.

Previous theoretical studies have addressed how synaptic pruning affects the properties of networks that store information in their connectivity matrix. Random pruning of synaptic connectivity (independent of synaptic strength) increases the efficiency of memory storage in Hopfield-type networks, i.e. the storage capacity in terms of numbers of stored attractors per available synapse increases with pruning (Sompolinsky 1986, Derrida, Gardner et al. 1987). Another line of work has characterized the statistics of synaptic connectivity in networks that optimize the robustness of fixed-point attractor states or sequences (Kohler and Widmaier 1991, Brunel, Hakim et al. 2004, Brunel 2016, Zhang, Zhang et al. 2019). In networks of excitatory and inhibitory neurons increasing the robustness of attractors led to networks with synaptic connectivity distributions that were sparse (i.e. many absent connections) but strong (i.e. the connections that remained were larger) (Kohler and Widmaier 1991, Brunel 2016, Zhang, Zhang et al. 2019). It was also shown that the measured distribution of synaptic strengths in cortex was consistent with the predictions of the theory (Brunel 2016). Thus, this work suggested that robust attractor dynamics lead to sparsely connected networks with fewer but stronger connections. Our work further extends these results by showing that pruning of weak connections in rate and spiking networks with temporally asymmetric Hebbian plasticity leads to large attractor basins for strong memories, while having minimal effects or eliminating memories that are weakly encoded.

Although we have reasonable network models for the computations that underlie cognition and learning (Sussillo 2014), like the models we have explored here, less is known about the specific neural systems in which these computations are implemented. For example, OCD is thought to reflect disordered computation in cortical-basal ganglia-thalamocortical circuits (Milad and Rauch 2012, Ahmari and Dougherty 2015, Sha, Versace et al. 2020). Whether OCD symptomatology reflects disordered computation in only one component of this circuitry, that is then propagated through the rest of the circuitry, or deficits in multiple nodes of the circuit, is unclear. Neurophysiology experiments have consistently shown that task variables related to affective learning have similar representations across nodes of CBTC circuits (Tang, Costa et al. 2022, Tang, Bartolo et al. 2024) and therefore circuit pathology may be distributed.

We have motivated our hypothesis from the perspective of strong memories that are embedded in networks before pruning, that subsequently come to dominate the attractor landscape after pruning. During adolescence, when weak synapses are pruned and stronger synapses are further strengthened (Chung, Wills et al. 2017, Li, Ma et al. 2017), emotionally intense memories come to dominate the attractor landscape, and less intense memories are pruned away. These dynamics can lead to the presence of unwanted, intrusive thoughts that are difficult to suppress. This can account for disorders including PTSD that follows childhood abuse and neglect, the effects of rumination in depression, early exposure to drugs of abuse in addiction (Anthony and Petronis 1995, Chambers, Taylor et al. 2003), and some forms of OCD in which the compulsions or rituals are linked to the obsessions in a straightforward way, like handwashing compulsions with contamination obsessions. This hypothesis can also link early life stress or adverse childhood events to the increase of the prevalence of psychiatric disorders as the early life stress can embed strong memories (McLaughlin, DeCross et al. 2019, Aloi, Crum et al. 2024) that become dominant through synaptic pruning. Studies of early life stress have also suggested delayed maturation of EEG dynamics that may reflect pruning (Andujar, Liuzzi et al. 2026), and a brain connectivity measure related to multiple psychiatric disorders correlates with delayed grey-matter volume reductions normally seen in adolescence (Xie, Xiang et al. 2023). Thus, there may be variations in developmental processes that further contribute to psychopathology.

It is not as immediate to account for OCD in which the compulsions do not have a reasonable link to the obsession. These cases may have overlap with delusional ideation (O’Dwyer and Marks 2000). It has been suggested that chaotic activity in dopamine neurons can lead to irrational associations (Kapur, Mizrahi et al. 2005). In effect, if the neuromodulator component of three-factor learning is dissociated from coherent events which normally give rise to intense emotions, bizarre associations could be formed and embedded in neural systems. Our hypothesis makes the strong claim that persistent ritualized behavior, or intrusive thoughts and memories that are difficult to suppress, are driven by strong attractor dynamics. And pruning and myelination leads to networks that are dominated by fewer but stronger attractors. The mechanism that gives rise to the strong embedded memories may in some cases be determined by coherent life events and in others by stochastic but strong activity in neuromodulatory systems that embed attractors that are only weakly associated with coherent life events, but nevertheless lead to strong patterns of activity in synaptic weights.

Ketamine and psilocybin, which are novel approved or under consideration treatments for depression and other psychiatric disorders, have been shown to increase spine density and therefore excitatory connections, in cortex (Savalia, Shao et al. 2021, Shao, Liao et al. 2021). Electroconvulsive therapy (ECT) may also increase spine density, because ECT induces seizures which could drive spine formation through activity dependent mechanisms. These treatments may alleviate clinical symptoms by increasing the capacity for plasticity and opening a window for other treatments like cognitive behavioral therapy, to work. However, it is also possible that increasing spine density may lead to clinical improvements by decreasing the strength of dynamical attractors. Adding random spines to pruned networks makes the sequential dynamics less robust (data not shown). As we have shown in the present paper, and in a previous study (Averbeck 2022), pruning increases the basin of attraction of dynamical attractors. While this can lead to improvements in cognition when the attractors represent beneficial patterns of activity, it can lead to disorders when the attractors represent negative emotional states.

## Methods

We used both rate-based and spiking neural networks to explore the hypothesis that pruning of weak connections can lead to expansion of dynamical attractors, especially those related to strong emotional memories. Here, we use the term ‘attractor’ in a loose sense, to refer to a sequence of network states that can be triggered from a large set of initial conditions. The implemented networks were developed from previous work, which explored in detail the sequence capacity of these models (Gillett, Pereira et al. 2020, Gillett and Brunel 2024).

## Rate-based networks

Rate-based neural networks were governed by the following equations:

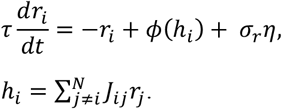

The variable *τ* is the time-constant of the single-cell dynamics (*τ* = .010 *s*), *h*_*i*_ is the synaptic drive to neuron *i, J*_*ij*_ is the connection weight from neuron *j* to neuron *i, N* is the number of neurons in the network (*N=*20000), *ϕ*(*h*) is the rate transfer function for single neurons, and *η* is a zero-mean white noise process. This transfer function was given by

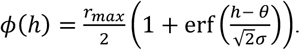

The variable *r*_*max*_ is the max firing rate of single neurons, *θ* is the input, *h* at which the firing rate is half its maximal value, and *σ* is the inverse gain or slope of the transfer function and erf is an abbreviation for the error function. For all rate network simulations, *θ* = 0.22, *σ* = 0.1, *r*_*max*_ = 1.

We used two different scenarios: One with a fixed connectivity matrix, assumed to have been structured by a past learning process; And the other where the connectivity matrix is built iteratively by a learning process. In the first scenario, connection weights in our networks are given by:

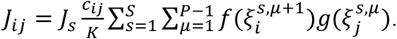

The variable *c*_*ij*_ indexes whether a connection exists (*c*_*ij*_ = 1) or not (*c*_*ij*_ = 0) from neuron *j* to neuron *i*. Networks were sparsely connected. Connections were defined randomly in the network by sampling from a binomial distribution with connection probability given by *p*. For rate simulations this was *p =*

0.01. *K* is the total number of connections to each neuron, S is the number of sequences, P is the length of each sequence, and 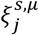 is the activity of neuron *j* in sequence *s* for element *µ*. The functions *f* and *g* can be used to implement thresholding. For all simulations in the rate network, these were the identity function. The patterns 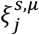 were randomly sampled from a standard normal distribution and were, therefore, approximately orthogonal to each other within and across sequences.

Sequence recall under noise-free conditions (no perturbation) was tested by setting *h* = *ξ*^*s*,1^. When perturbations were applied, the network was initiated with *r* = *ϕ*( *ξ*^*s*,1^) + *σ*_*p*_*η*. The value of *σ*_*p*_ is given in the relevant figures. Correlations between activity in the network and sequences were calculated as:

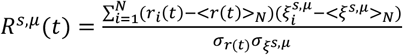

For the acquisition and extinction learning simulations, synaptic weights for each sequence are updated at each iteration according to

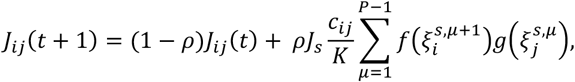

where *J*_*s*_ is the maximum connection strength. This implements the fact that plasticity saturates (Nguyen-Vu, Zhao et al. 2017). During extinction learning a new sequence is learned that links the cue to no outcome, and the previous sequence that links the cue to the aversive outcome is forgotten. This is done by setting *J*_*s*_ = 0 for the sequence that is being extinguished, because it is not being experienced. Extinction is slower than acquisition (Quirk 2002), and therefore we used a lower learning rate for extinction. Thus, during extinction and reacquisition, we end up with a superposition of sequences, given by

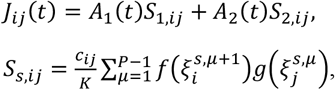

where *A*_*s*_ indicates the accumulated synaptic strength associated with each sequence.

Three-factor learning rules modulate the strength of synaptic connectivity, driven by Hebbian plasticity, including STDP (Fremaux and Gerstner 2015, Kusmierz, Isomura et al. 2017). These are called three-factor rules, because they require specific timing between a pre-synaptic spike, a post-synaptic spike, and the presence of a neuromodulator. In the absence of the neuromodulator the plasticity does not take place, and the concentration of the neuromodulator may modulate the strength of the plasticity. Memory for events that trigger stress responses engage the glucocorticoid (cortisol) and noradrenergic systems (Shields, Sazma et al. 2017, Shields, Hunter et al. 2022), which may modulate synaptic plasticity via three-factor learning rules (Shouval and Kirkwood 2025). In our networks, when we embedded multiple sequences of different strengths, this was implemented by assuming both the learning rate, *ρ* and the maximum synaptic strength *J*_*s*_ were modulated by a mono-amine neurotransmitter, for example dopamine or norepinephrine (Reynolds, Hyland et al. 2001). In this case, both were functions of the dopamine concentration, which was assumed to be a function of the emotional intensity of the event. In the present results we only explored the implications of this scaling for the maximum strength of the memory, however, it would be possible to expand this to consider other aspects of this scaling.

Synaptic pruning was implemented by rank ordering the absolute value of all synaptic connections present in the network (i.e. *c*_*ij*_= 1) and setting a fraction of the lowest rank synapses to 0, (*c*_*ij*_ = 0).

Specifically, we sorted the synapses such that 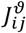 was defined such that 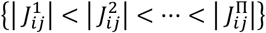 where ∏ is the total number of synapses in the network. We then set all synapses 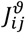 to 0, for *ϑ* ≤ *p*_*f*_ ∏, and *p*_*f*_ was the fraction of synapses to prune. The remaining synapses were then reweighted, such that the total synaptic strength *w* in the network was constant before and after pruning. The average input to each neuron is, therefore, kept constant, consistent with previous theory work (Kohler and Widmaier 1991, Brunel 2016, Zhang, Zhang et al. 2019, Iatropoulos, Gerstner et al. 2025):

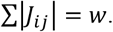

### Spiking networks

We simulated networks of leaky integrate and fire (LIF) spiking neurons. We first simulated a single network of 30,000 neurons to examine sequence recall in a single network. We then simulated two coupled E-I networks with 15,000 neurons each. The number of inhibitory neurons, N^I^, was always 0.25*N^E^. The membrane update equations for the excitatory and inhibitory neurons are given by:

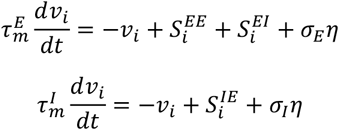

The variable *η* was a zero mean gaussian white noise process, and *σ*_*E*_ and *σ*_*E*_ were the standard deviations of the excitatory and inhibitory noise, respectively. The post-synaptic potential update equations are given by

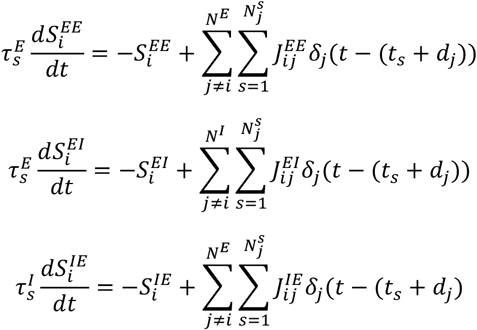

Where the variable *d*_*i*_ was the synaptic delay associated with pre-synaptic neuron *j* and *N*^*s*^ is the number of spikes in neuron *j*. Local delays were always 1 ms. Delays between populations depended on the synaptic delay distribution and are shown in the figures. Each neuron was assumed to have a fixed synaptic delay to all neurons in the population to which it projected, where the delay was given by that neuron’s level of myelination, axonal caliber and distance between nodes of Ranvier (Waxman 1980).

Note that all inhibitory connections were local and therefore all inhibitory synaptic delays were 1 ms. Neuron *i* fired a spike at time *d*_*s*_ if *v*_*i*_ > *V* _*threshold*_. After firing a spike, the membrane potential was reset to *V*_*rest*_. Furthermore, neurons had a refractory period and were held at their resting membrane potential, *V*_*rest*_ for 1 ms after firing a spike. Additionally, membrane potentials for excitatory neurons were not allowed to go below the inhibitory reversal potential, which was 10 mV below the reset potential.

Synaptic weights, *J*_*ij*_, were constructed using the approach outlined previously (Gillett, Pereira et al. 2020, Gillett and Brunel 2024). All structured connectivity was implemented between excitatory neurons. Inhibitory neuron synaptic weights were fixed for all connections. Inhibition and excitation have to be balanced to maintain firing rates within a reasonable range. Excitatory synaptic weights were defined by two steps. First, weights were structured using the same connectivity rule used for rate networks, except incorporating synaptic delays,

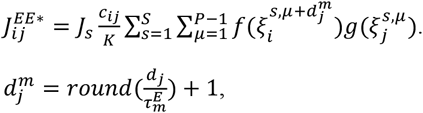

where the round function rounds its argument to the nearest integer. For spiking networks, the function *f* and *g* were given by:

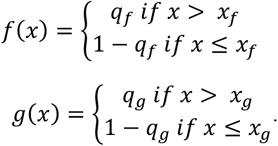

The variable *c*_*ij*_ indicates the presence of a connection (*c*_*ij*_ = 1) or not (*c*_*ij*_ = 0). The probability of a connection between excitatory neurons (*p*^*EE*^), inhibitory and excitatory neurons (*p*^*EI*^), and excitatory and inhibitory neurons (*p*^*IE*^) within a local population, as well as the connection between excitatory neurons between populations 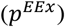 are given in Table 1. The delay was expressed in units of the membrane time constant, because the sequences evolved on a time-scale set by the membrane time constant. Note that all local connections had 1 ms delays, and therefore the synaptic connectivity in local networks was not affected by myelination. Only long-range connections between areas were affected by myelination. It is difficult to constrain the distribution of spike time delays that would be expected in the human brain. In the mouse, two studies have shown that the distribution of delays depends on myelination (Salami, Itami et al. 2003, Kato, Wake et al. 2020). These studies suggest that some disruption in myelin can lead to standard deviations of delays on the order of several milliseconds, in thalamocortical circuits. Given the massive scaling of distance from the mouse to the human brain, it seems likely that developmental effects may lead to distributions on the order of 10s of milliseconds. We have explored this range in the results.

**Table 1.** Parameter values used in spiking network simulations.

| Parameter | Value |
|---|---|
| J<sup>S</sup> | 6.3 |
| *τ*<sub>*m*</sub><sup>*E*</sup> | 0.010 s |
| *τ*<sub>*s*</sub><sup>*E*</sup> | 0.020 s |
| *V*<sub>*threshold*</sub> | 20 mv |
| *V*<sub>*reset*</sub>, *V*<sub>*rest*</sub> | 0 mv |
| *V*<sub>*reversal*</sub><sup>*E*</sup> | -10 mv |
| τ<sup>I</sup><sub>m</sub> | .002 s |
| τ<sup>I</sup><sub>s</sub> | .005 s |
| σ<sup>E</sup> | 5.0 |
| σ<sup>I</sup> | 5.0 |
| *p*<sup>*EE*</sup> | 0.08 |
| *p*<sup>*EI*</sup> | 0.04 |
| *p*<sup>*IE*</sup> | 0.04 |
| *p*<sup>*EEx*</sup> | 0.04 |
| *x*<sub>*f*</sub>, *x*<sub>*g*</sub> | 1.5 |
| *q*<sub>*f*</sub> | 0.8 |
| *q*<sub>*g*</sub> | 0.932 |

Weights were then passed through a threshold linear (relu) nonlinearity, such that only positive weights remained,

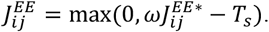

For all simulations, *ω* = 1.0, *T*_*s*_ = −0.0038. The connections to the inhibitory population, and from the inhibitory population to the excitatory population were fixed for all neurons (i.e. 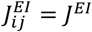 for all *i,j*), and were set using the mean-field approach outlined in (Gillett, Pereira et al. 2020). Note that EI connections from inhibitory to excitatory neurons were negative. All other connections weights were positive. Networks were sensitive to inhibition levels such that when inhibition was too high neural activity dropped to low levels and sequence recall failed and when inhibition was too low neural activity diverged and sequence recall failed. Inhibition was not adjusted over pruning as excitatory drive was rescaled and therefore the total excitatory drive was constant, and therefore the amount of inhibition necessary to balance the excitation was also constant, in the network.

For sequence recall in spiking networks, activity was initiated by setting 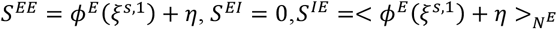. The variable *η* is Gaussian white noise added to the initial activity to perturb it away from alignment with the first sequence element. When activity was started in only one population, *ξ*^*s*,1^ was set to zero for all neurons in population 2, for sequence initiation.

When we pruned synapses in the spiking networks, we pruned only the excitatory synapses. The same procedure was used. We pruned a fraction of the weakest excitatory synapses and then reweighted the remaining synapses such that the total excitatory drive in the network was kept constant. Inhibition was not changed.

For the decoding in the acquisition and extinction experiment, we decoded the representation of the last sequence element. We first calculated the minimum Euclidean distance between the network activity throughout recall and the last element for both the aversive and safety sequences,

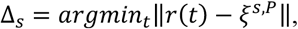

where s indexed either the aversive or safety sequence. This was then passed through a softmax nonlinearity to normalize values between 0 and 1,

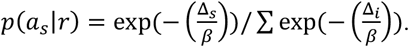

The variable *β* was set to 10. This scales the slope of the transitions.

## Data and Code Availability

There is no data associated with this manuscript. The code for the manuscript is available on the first authors GitHub page, https://github.com/baverbeck/Pruning-RNNs.

## Author Contributions

B.B.A and N.B. conceived the study. B.B.A. wrote the code and authored the manuscript. N.B edited the manuscript.

## Acknowledgements

This work was supported by the intramural research program of NIMH (ZIA MH002928). Simulations were carried out on the NIH/HPC Biowulf cluster (http://hpc.nih.gov). The contributions of the NIH authors were made as part of their official duties as NIH federal employees, are in compliance with agency policy requirements, and are considered Works of the United States Government. However, the findings and conclusions presented in this paper are those of the authors and do not necessarily reflect the views of the NIH or the U.S. Department of Health and Human Services.

